# Bleaching-resistant, near-continuous single-molecule fluorescence and FRET based on fluorogenic and transient DNA binding

**DOI:** 10.1101/2022.04.19.488730

**Authors:** Mirjam Kümmerlin, Abhishek Mazumder, Achillefs N. Kapanidis

## Abstract

Photobleaching of fluorescent probes limits the observation span of typical single-molecule fluorescence measurements and hinders observation of dynamics at long timescales. Here, we present a general strategy to circumvent photobleaching by replenishing fluorescent probes via transient binding of fluorogenic DNAs to complementary DNA strands attached to a target molecule. Our strategy allows observation of near-continuous single-molecule fluorescence for more than an hour, a timescale two orders of magnitude longer than the typical photobleaching time of single fluorophores under our conditions. Using two orthogonal sequences, we show that our method is adaptable to Förster Resonance Energy Transfer (FRET) and that can be used to study the conformational dynamics of dynamic structures, such as DNA Holliday junctions, for extended periods. By adjusting the temporal resolution and observation span, our approach should enable capturing the conformational dynamics of proteins and nucleic acids over a wide range of timescales.

## Introduction

Single-molecule methods have transformed the study of biological systems by enabling detailed interrogation of the structure, dynamics, and function of individual molecules. These methods offer unique insight into many biological molecules and processes, including the folding of proteins, the mechanisms of gene expression and maintenance, the structurefunction relationships of molecular assemblies, and the coupling of large macromolecular machines, both in vitro and in vivo.^[1]^

In particular, single-molecule fluorescence (SMF) spectroscopy and microscopy studies have been very popular, since they are sensitive, versatile, and compatible with use in living cells.^[2]^ Some SMF studies involve labelling the biomolecule of interest with a single fluorophore^[3]^, which allows detection of the labelled molecule, and localisation of its position with high precision. These capabilities in turn enable measurements of molecular stoichiometries, as well as super-resolution imaging and single-molecule tracking.^[4–10]^ Other SMF studies involve single-molecule FRET (smFRET), which typically uses two complementary fluorophores to monitor distances in the 2-10 nm range and can be used as a molecular ruler^[11,12]^. smFRET can also report on the kinetics of conformational changes and relative motions of interacting species, which helps to deduce the sequence of events in many biological processes.^[13]^

Most SMF (and more generally, fluorescence microscopy) methods, however, are still severely limited by photobleaching, which is the irreversible photo-destruction of the fluorescent probes (either organic dyes or fluorescent proteins) used to label the biomolecules of interest; such limitations have been evident since the early days of SMF.^[14,15]^ Due to photobleaching, the photon budget per fluorophore (i.e., the number of photons emitted before the end of the observation) remains limited, and restricts observation time to the low-minute timescale.^[16,17]^This limitation remains despite significant improvement through the use of fluorescence stabilisation systems (such as oxygen scavengers and triplet-state quenchers)^[18]^.

A possible way to overcome photobleaching is to exchange the fluorescent labels during an ongoing experiment. This can be achieved using transient or reversible interactions between the target molecule under study and the fluorophore, provided that the system allows for an exchange of an attached fluorescent probe with a new probe *before* the attached probe gets photobleached.

Exchanging bleached fluorescent labels with fresh ones has been explored for self-healing and regeneration of DNA nanostructures,^[19]^ where an incubation with fresh “staple” strands repaired the effects of photo-damage to fluorescently labelled staples. Further, transiently binding fluorophores have been used to study targets for extended periods, especially in single-molecule localisation microscopy (SMLM), with an early example being the method of Points Accumulation for Imaging in Nanoscale Topography (PAINT).^[20]^ Such transient binding was also used in DNA-PAINT, where short labelled DNA strands (“imagers”) bind to complementary “docking strands” on target biomolecules such as DNA nanostructures^[21]^and proteins.^[22]^ Exchanging fluorescent labels has also been combined with stimulated emission depletion (STED) microscopy to enhance photo-stability.^[23]^

To achieve super-resolution, methods like DNA-PAINT require temporal separation of single-molecule fluorescence signals from a diffraction-limited area and thus need extensive “dark intervals” (i.e., during which the target is not bound by a transient label, and thus is not fluorescent). Consequently, these methods cannot provide the *continuous* signal needed to monitor the presence or motions of a molecular target over extended observation spans. In principle, the dark interval between the binding of two transient labels to the same target can be decreased by increasing the rate of binding, either by increasing the transient label concentration, and/or changing the properties of the transient label to increase the on-rate constant. However, the concentration of fluorescent transient labels cannot be increased much above 30 nM, as unbound labels contribute to the fluorescence background and degrade the SNR of the measurement^[24–27]^.

One approach to generate continuous fluorescence over long time spans is to multiplex binding sites for transient DNA labels and optimise binding/unbinding kinetics, which can allow for single-particle tracking for hours.^[28]^ Whilst this approach is attractive for SMF measurements with a single fluorophore, it is incompatible with smFRET studies, where the multiple fluorophores along the docking DNA strand will create an uninterpretable web of fluctuating photophysical interactions between many potential FRET donor and acceptors. A recent approach built on the concept of dye-cycling by implementing it on a *single* binding site, thus enabling extended smFRET experiments^[28]^ on Holliday junctions (HJ) using the reversible binding of fluorescently labelled 9-nt-long ssDNAs carrying either a FRET donor and a FRET acceptor (“cyclers”)^[29]^. However, despite measuring in an SNR regime that did not allow resolution of the dynamics of the HJ, the maximum concentration of transient fluorescent labels was limiting the temporal sampling to only ~50% per cycler, and thus having single-molecule targets spending only ~25% of time in a doubly-labelled state, which is a prerequisite for FRET measurements.^[29]^

The limitations of the past studies vividly highlighted that the fluorescence background due to unbound labels is the main bottleneck to achieving continuous fluorescence traces.

Here, we address these limitations by introducing a transient binding approach that optimises background suppression and label exchange to enable near-continuous, bleaching-free single-molecule fluorescence observations via **R**enewable **E**mission via **F**luorogenic and **R**epeated **s**sDNA**H**ybridisation (REFRESH, Figure 1A). We also extend this approach to smFRET (REFRESH-FRET, Figure 1B). The target biomolecule is modified with a short ssDNA docking strand which is recognised by a labelled complementary DNA (a “renewable label” or “r-label”). Importantly, our approach involves the design of fluorogenic r-labels that enable measurements in high concentration regimes which allow for continuous (or near-continuous) emission of fluorescence from a target biomolecule with both high temporal resolution *and* extended observation span (>1 hr, Figure 1A, bottom). We show that we can specifically label two sites within a molecule and enable smFRET measurements over the same period. Finally, we show that transient labelling is fully compatible with dynamic biomolecules by monitoring the conformational dynamics of HJs using long-lived smFRET measurements. Our strategy can be easily tuned to adapt its temporal resolution and observation span to a plethora of biological systems and applications.

**Figure 1.**
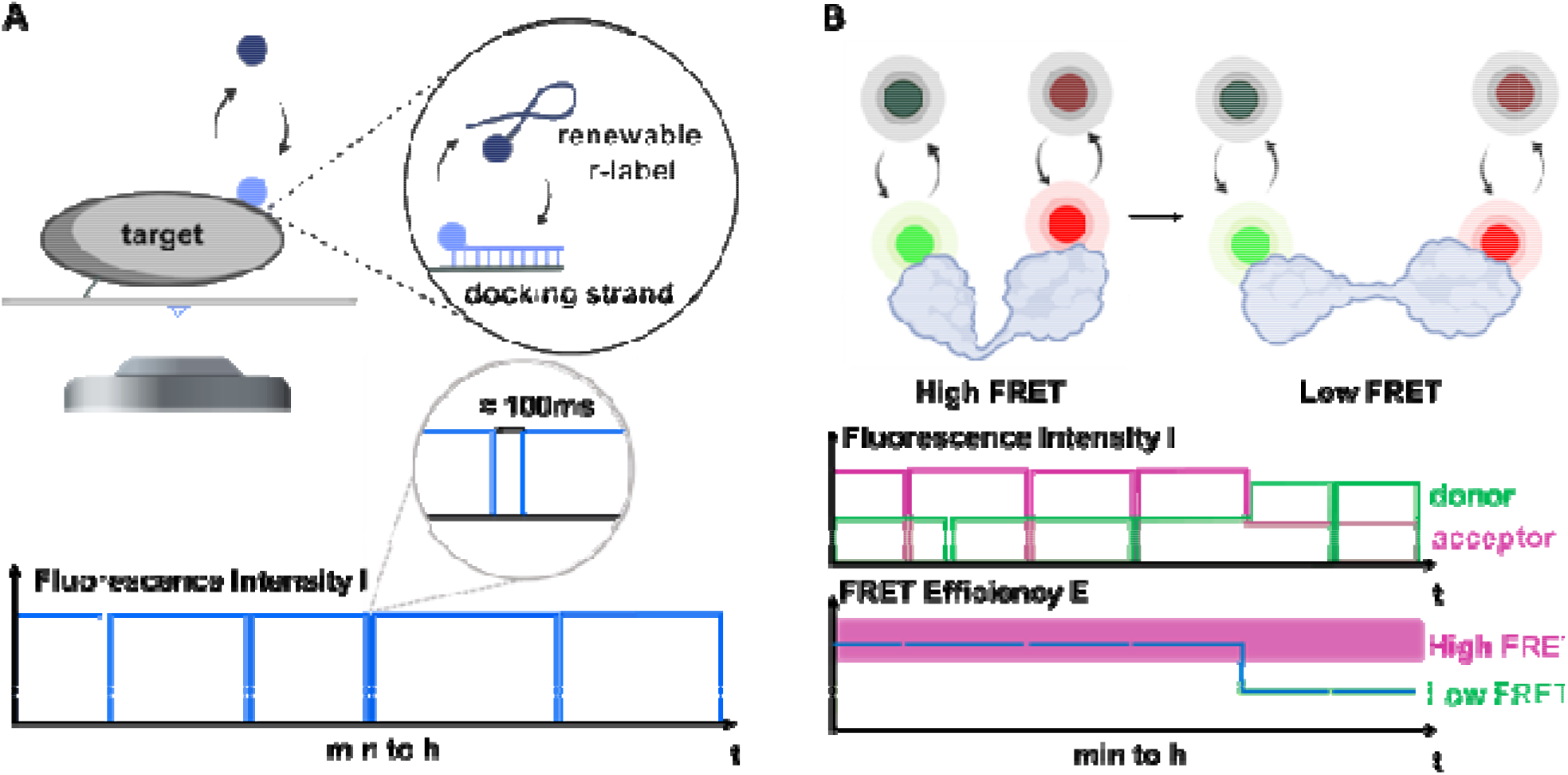
The principle of REFRESH. A. Continuous exchange is facilitated by transient hybridisation of short, fluorogenicssDNA probes (r-labels) to complementary docking strands bound to a target molecule of interest (top). The schematic fluorescence trace (bottom) shows how, if intervals of exchange are short (below or on the order of the frame rate), a near-continuous fluorescence signal can be observed. B. REFRESH-FRET: The same labelling strategy is now applied to donor and acceptor dyes attached to a target of interest which undergoes conformational changes. The observed donor and acceptor traces (middle) can be used to calculate FRET efficiencies (bottom) and monitor conformational dynamics.

## Results and Discussion

### Design Principles for REFRESH

Since the labelling of a target biomolecule (hereafter, the “target”) is based on a series of reversible binding events, the resulting time-traces from a target will contain dark intervals; the ideal traces should have as few and as short dark intervals as possible, achieving a temporal target sampling that approaches 100%. This aspiration for near-complete sampling creates two requirements: first, dark intervals due to r-label-exchange events (where a dissociated r-label is replaced by a new one) need to be short, and ideally should occur on a timescale similar (or shorter) than the exposure time of the single-molecule imaging experiment (typically, in the 20-200 ms range). Second, dark intervals due to any bleaching of an r-label while bound to the target need to be minimised both in number and duration. To fulfil these requirements, we use the following set of strategies: (I) To achieve high on-rates, we optimise DNA sequences by avoiding intramolecular complementarity. (II) To minimize dark intervals, we employ high r-label concentrations, a condition facilitated by r-labels that are fluorogenic, i.e., when unbound, remain in a dark, quenched state. (III) To increase the photobleaching lifetime of the fluorophore, we use a photo-stabilisation system. (IV) To ensure r-label dissociation *before* bleaching, we tune the off-rate of the r-label from the target. (V) To use the photon budget of each r-label efficiently, we maximise the binding time between r-label and target within the limit set by photobleaching.

### Sequence Selection

The DNA-sequences used for the r-labels were chosen as follows: for the initial sequence (red r-label), we selected a DNA length of 10-12 nt to provide a “bound time” that is long enough to ensure efficient use of the photon budget of the fluorophores, and short enough to avoid photo-bleaching in the bound state (which, for a single ATTO647N fluorophore, is described by k_bleach_ = 0.003 s^−1^ under continuous excitation by a laser power of 1.4 mW at 640 nm). Broadly speaking, k_off_ has to be 10-100 fold larger than k_bleach_, and k_on_ should be 10-100-fold larger than k_off_. Importantly, our r-labels are considerably longer than the imager strands employed in super-resolution techniques, since we want to minimise dark intervals. The ultimate goal of a high on-rate (which minimises dark periods) could be facilitated by a high r-label concentration, but could also be influenced by the sequence: repetitiveness in sequence (e.g., consistent of repeats of a three-base motif) combined with a longer sequence have been shown to increase the on-rate.^[30]^ We also avoided interactions within the sequence by choosing just two non-complementary bases per sequence (e.g., only thymine (T) and guanine (G)), and calculated ΔG values from Santa Lucia et al.^[31]^, and on- and off-rates for DNA hybridisation using an algorithm by Zhang et al.^[32]^

Using these design principles and experimental testing, we identified a first suitable DNA r-label, which featured an 11-nt long sequence with a low guanine/cytosine (G/C) content (3 out of 11); such an imager DNA strand enables fluorogenicity (see next section), while dehybridising faster than a strand with the same length but a high G/C content. Specifically, the red r-label showed a mean t_off_ of ~3.3 s (or k_on_ ~ 0.30 s^−1^) and a mean t_on_ of ~15 s (or k_off_ ~ 0.07 s^−1^) at 20 nM.

For the second r-label sequence, we built on the fluorogenic DNA-PAINT imager sequence design in Chung et al., which extended the imager length to 15 nt and used mismatches between imager and docking strand to reduce bound times and allow for blinking and superresolution imaging.^[33]^Starting with their green imager sequence, we introduced a higher degree of complementarity and finally selected the most suitable sequence out of seven experimentally tested ones for REFRESH (with a mean t_off_ of ~12 s (or k_on_ ~ 0.08 s^−1^) and a mean t_on_ of ~22 s (or k_off_ ~ 0.05 s^−1^) at 100 nM. (For simplicity, we have only stated mean values for the dwell times here. Further characterisation of the hybridisation kinetics of both r-labels can be found in the supplementary material and Figure S1.)

### Fluorogenic Strategy

To keep the dark intervals low, our technique relies on using high concentrations of unbound labels (100 nM – 1 μM); this, in turn, leads to a significant fluorescence background that needs to be suppressed. A standard way to reduce the fluorescence background is using an evanescent excitation field in a total-internal-reflection fluorescence (TIRF) microscope; however, this mode of microscope still cannot allow detection of single immobilised molecules in the presence of >50 nM of unbound label.^[24–27]^ To further suppress the fluorescence background, we considered a fluorogenic strategy that quenches the fluorescence of unbound labels, but allows for substantial recovery of fluorescence upon label binding.

For a red label, we use a short ssDNA labelled with two ATTO647N fluorophores (one on either end) that exhibit contact-mediated quenching in solution (see SI and Figure S2); when bound to the target, the state of quenching is lifted, leading to the appearance of fluorescence corresponding to two ATTO647N fluorophores.^[24]^ Use of this fluorogenic strategy improves the SNR by a factor of 4 and effectively makes the r-label (as a unit) more photostable, since complete loss of fluorescence requires bleaching of both fluorophores and thus will require more time to occur.

For the green label, we have used r-labels that contain a pair of a green fluorophore (Cy3B) and a dark quencher (DQ), which serves as a non-fluorescent FRET acceptor to the Cy3B fluorescence in the unbound r-label. To allow for sufficient fluorogenicity in the bound state, we use an extended r-label length of 15 nt with mismatches between r-label and docking strand, as was done recently for fluorogenic DNA-PAINT imagers.^[33]^ This allowed for a gain in signal-to-noise ratio (SNR) by a factor of 16 (see SI, Figure S2), and allowed for imaging of single-molecule binding sites using a TIRF microscope with up to 5 μM r-label (see Figure S3).

### Near-continuous SMF with REFRESH

We first implemented our renewable strategy on a surface-immobilised DNA target containing a docking DNA strand complementary to our respective r-label (note: the target is the HJ used later for smFRET experiments). The target was also labelled with a Cy3B fluorophore, which served as a localisation signal. During the experiment, we first localised our target using the green emission channel (Figure 2A, left); we then added the r-label strands, and recorded movies under red or green excitation (Figure 2A, right), which were then used to generate time-traces.

**Figure 2.**
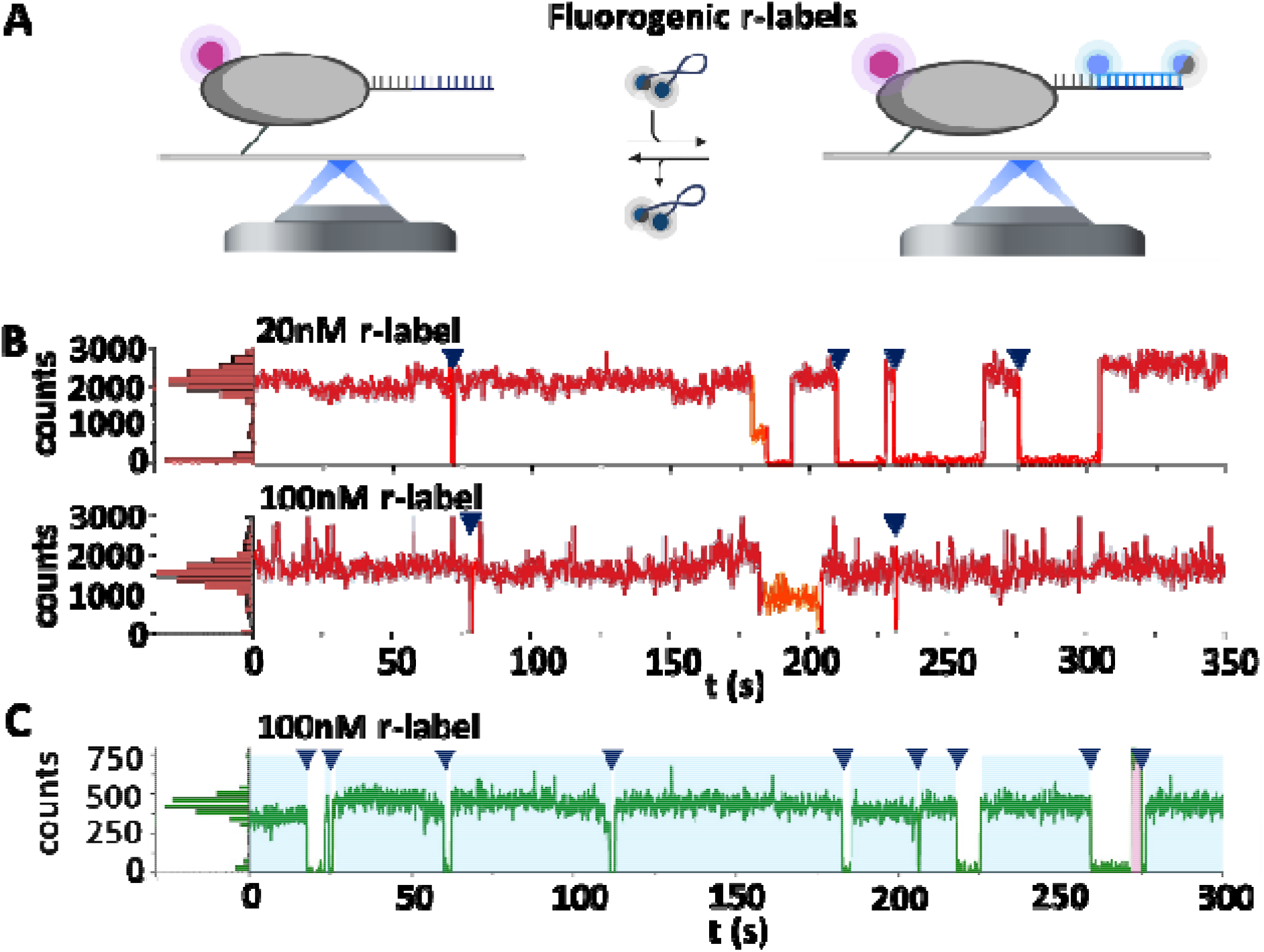
REFRESH allows for continuous single-molecule fluorescence observations: A. The target molecule is localised on the surface via a localisation label (magenta), observed in an emission channel different from that of the r-labels. After the addition of r-labels (carrying two ATTO647N or Cy3B/BHQ2), binding and unbinding can be observed at co-localising spots. B. Example traces at 20 nM (top) and 100 nM (bottom) red r-label. C. Example traces of 100 nM green r-label. Blue shaded areas: bound intervals of the complete r-label; triangles indicate de-hybridisation; yellow shaded areas: interval with only one emitting ATTO647N; magenta shaded area: interval without functional BHQ2. For more traces of binding of the green and red r-label, see Figures S5 and S6, respectively.

We generated time-traces at different concentrations of the red r-label (Figure 2B). The fluorescence traces showed several intervals of high intensity (≈2,000 counts), corresponding to an r-label being bound to the target (blue shading) followed by disappearance of the fluorescence signal (blue triangles), which we attribute to r-label dissociation (and *not* to bleaching, which would have led to a step-wise decrease in intensity due to the presence of two fluorophores per r-label). As expected, the dark intervals became shorter with increasing r-label concentration (compare Figure 2B and C), and for concentrations of >100 nM, become negligible (<2 %). Occasionally, bleaching of one fluorophore occurs, reducing the fluorescence signal by ~50 %, to ≈1,000 counts (yellow shading), which also suggests that de-quenching upon target binding is complete, with no significant impact of any remaining contact-mediated quenching or homo-FRET processes on the quantum yield. Further decrease to baseline intensity is attributed to r-label unbinding or bleaching of the second fluorophore.

We performed similar experiments using a target carrying the docking strand for the green r-label. To avoid FRET interactions between localisation signal and r-label, we used a second Cy3B dye as localisation signal which was bleached before addition of the r-labels. Our traces at 100 nM green r-label (Figure 2C) show clear r-label binding events, detected as an increase in signal from background level to ≈450 counts (blue shading). We observe a mean bound time of ~22 s, which indicates a turnover faster than for the red r-label (with any dwells shorter than the 100-ms frame time being inaccessible).

Occasionally, we observe a signal level of ≈800 counts (magenta shading), which we attribute to an r-label without functional quencher, which thus appears brighter due to the lack of FRET. The observed increased signal in the absence of quenching is consistent with expectations from the 14-bp separation between fluorophore and quencher (~5.5 nm) and the Förster radius between Cy3B and BHQ2(~6.1 nm).^[34]^We also see occasional very short spikes of the same level of fluorescence during binding events, which we attribute to quencher blinking and a transient absence of the FRET quenching process.^[35]^ At 100 nM, we observe a temporal sampling of ~70%. To increase the temporal sampling further, we can increase the r-label concentration to e.g., 300nM for the following smFRET experiments.

Our results clearly established that we can perform continuous observations on immobile biomolecules using renewable labelling based on the hybridisation of short, ssDNA labels. Importantly, due to constant r-label exchange, the trace length was not limited by the bleaching of individual fluorophores, and since the fluorescence signal was lost for very few frames at a time, we achieved a very high temporal target sampling, which reached up to 98%.

### Observation of Conformational Dynamics for hours using REFRESH-FRET

We then moved to the experimental implementation of smFRET measurements using r-labels. As a model system, we chose a HJ, a well-studied dynamic four-arm DNA structure that allows us to monitor repeated interconversions between two conformational states distinguishable using FRET.^[36]^

We first assembled a “standard” HJ by using four 22 nucleotide-long ssDNA strands, one of which carries a covalently attached FRET donor and a second carrying a covalently attached FRET acceptor; this labelling strategy results in the fluorophores appearing at the ends of two of the HJ’s four arms (Figure 3A). In the two main HJ conformational states, the fluorophores are positioned at very different distances from each other, resulting in two distinct FRET efficiencies (E, a high FRET state of E ≈ 0.75 and a low FRET state of E ≈ 0.25).

**Figure 3.**
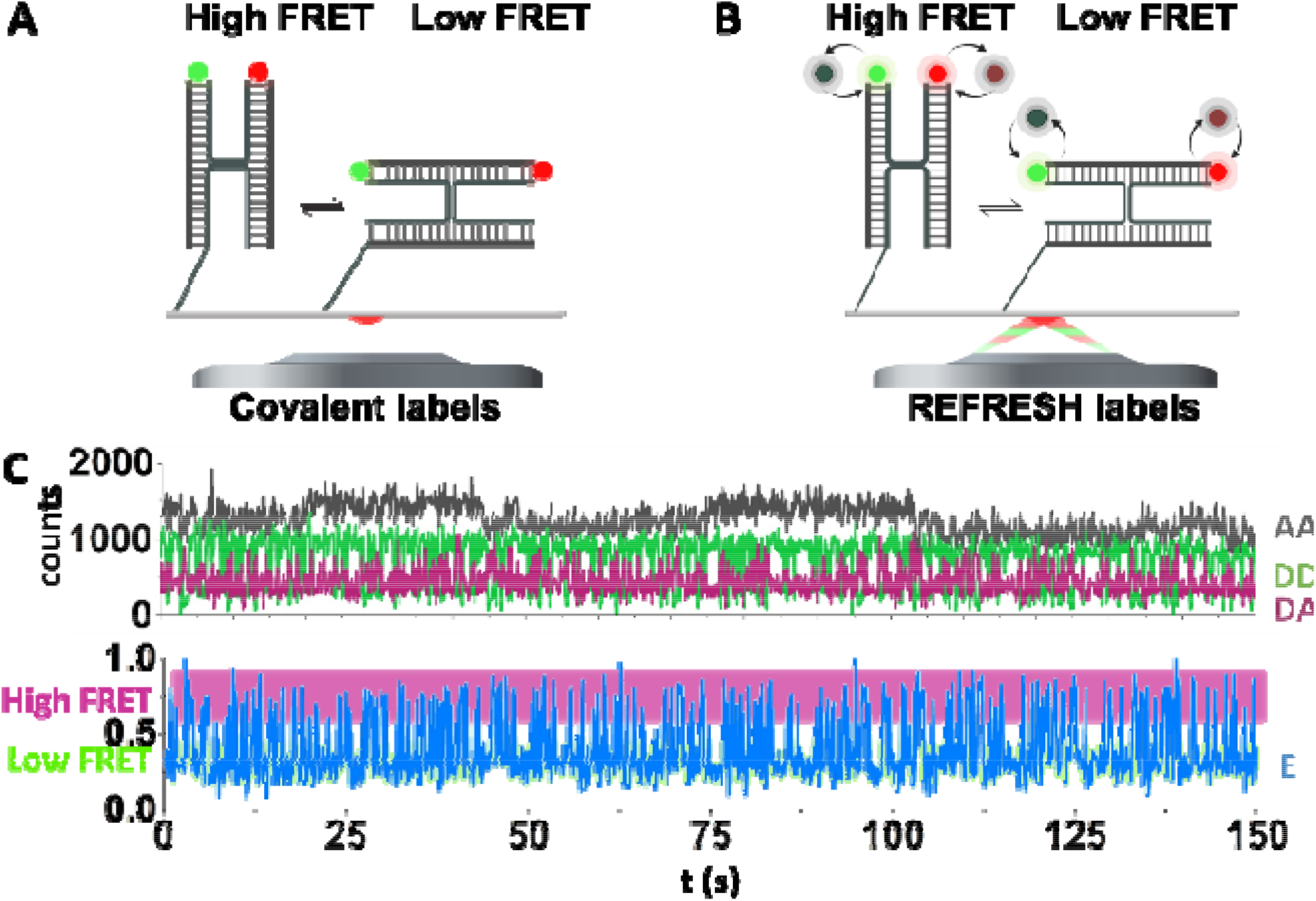
Observation of conformational dynamics using REFRESH-FRET. A. Reference structure of a covalently labelled HJ. B. HJ with exchanging acceptor labels: The X- and R-strand of the HJ carry extensions which serve as specific binding site for the fluorogenic r-labels. C. Intensity-vs-time trace of the reference HJ with AA, DA, and DD signal (top), from which the apparent FRET efficiency (E) was calculated. The anti-correlated fluctuations in DD and DA and the fluctuations in E indicated FRET dynamics between a high-FRET state (E ≈ 0.75) and a low-FRET state (E ≈ 0.25).

To monitor both FRET and fluorophore stoichiometries, we used alternating-laser excitation (ALEX) of the immobilised molecules using 200-ms frame times (100-ms exposure/channel). For each ALEX frame (see *Methods*), the AA (emission in the acceptor channel during acceptor excitation) signal reports on the presence of the acceptor, and the DD and DA (emission in the donor or acceptor channel during donor excitation, respectively) signals are used to observe FRET and conformational changes. Figure 3B shows a representative example of a fluorescence trace recorded from the reference HJ: the AA trace shows an intensity of ~1,500 counts, and features slow fluctuations between two spectral states of ATTO647N, which have been described before.^[37]^ On the other hand, DD and DA show anti-correlated fluctuations indicating dynamic FRET processes, which are also clearly reflected in the apparent E trace (bottom panel), which shows transitions between high (E ≈ 0.75) and low (E ≈ 0.25) values.

To implement REFRESH-FRET, we extended the strands carrying the reporter dyes by docking sequences complementary to our green and/or red r-labels. In solution, we provide the required fluorogenic r-labels which bind specifically to their docking strands (Figure 3A). In the two main HJ conformational states, the fluorophores are again positioned at different distances from each other, resulting in two distinct FRET efficiencies. Since the observation time of the REFRESH-HJ is essentially immune to the photobleaching, we can monitor it for extraordinarily long time-spans; we thus recorded continuous traces using 100 nM of red and 300 nM of green r-label, and an exposure time of 100 ms/channel/frame (Figure 4) for one hour, several orders of magnitude longer than the bleaching time of the individual fluorophores. We replaced the buffer in the chamber continuously (with a full volume exchange every 5 min) to replenish the stock of r-labels and the photostabilisation system.

**Figure 2:**
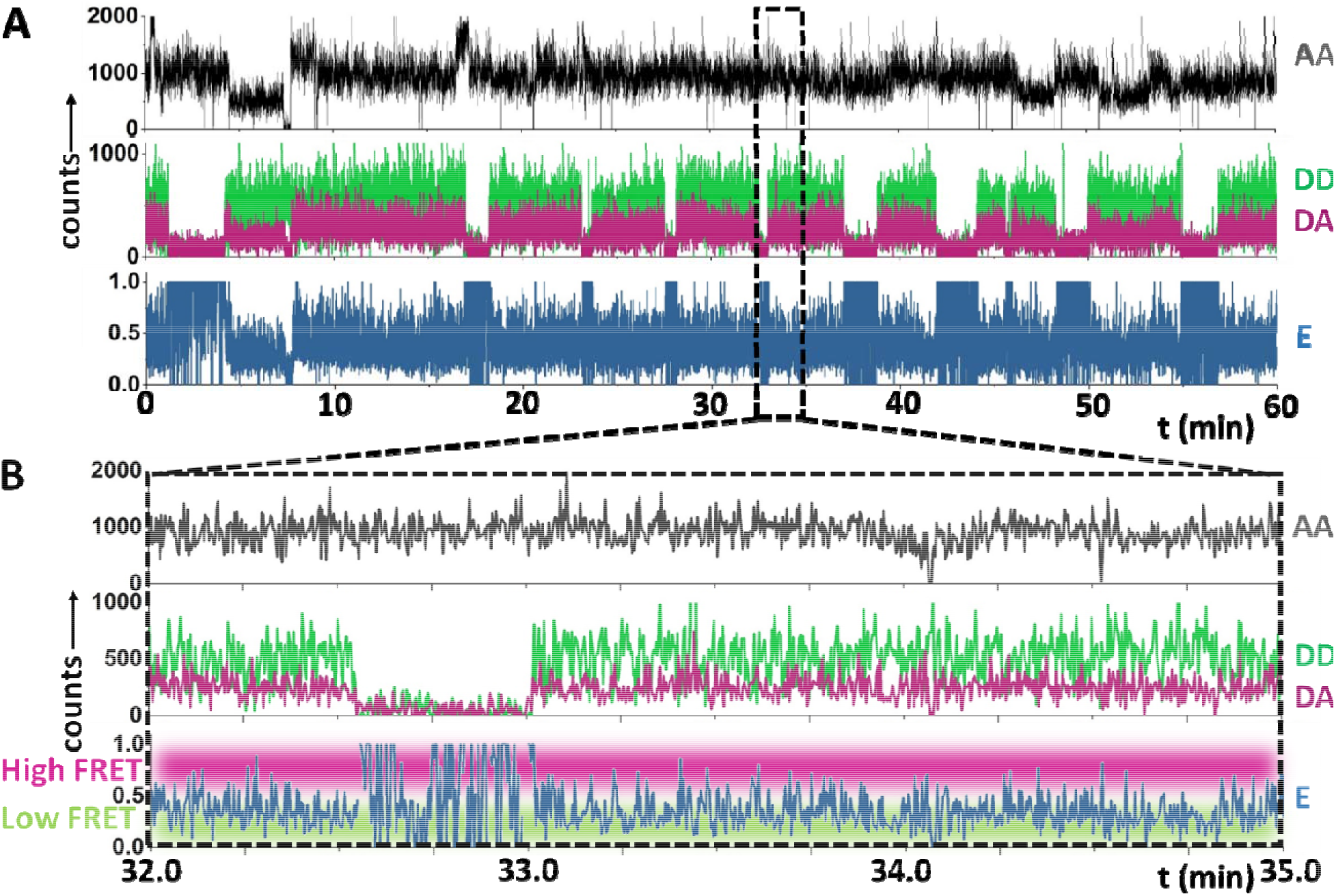
Representative traces from the REFRESH-HJ using 100 nM red and 300 nM green r-label. A. Full trace with AA (top), DD, and DA channel (middle) and calculated FRET efficiency (E, bottom). B. Zoom-in of the trace in A. The anti-correlated fluctuations in the DD and DA channels (middle panels) and the fluctuating E trace (bottom) indicate a dynamic interchange between a high-FRET state (E ≈ 0.75) and a low-FRET state (E ≈ 0.25). Further examples of FRET traces can be found in Figure S7.

Upon ALEX excitation, the AA trace specifically reports on the presence of the red r-label (by monitoring directly the acceptor presence and emission status), whilst the DD trace reports on the green r-label (by monitoring the donor presence and emission status) and the FRET efficiency. The presence of a significant DA intensity is only expected when both dyes are present at the same time and will report on the conformational state of the HJ. The observed time traces show the same pattern of binding events as shorter traces in Figure 2, with the additional feature that, whenever r-labels bind simultaneously, the same anticorrelated dynamics in the DD and DA intensity as with the reference HJ can be observed (zoomed-in segment, Figure 4B). The fluorophore fluctuations indicate FRET dynamics, with the E value showing clear transitions between high (E ≈ 0.75) and low (E ≈ 0.25) values. The FRET efficiencies for the two states are similar to the reference HJ, which validates the choice of the fluorophore location on the r-labels.

Inspection of the traces shows that the AA signal is near-continuous over the recorded period at an r-label concentration of 100 nM, whilst the DD trace shows still periods of time without fluorescence (~20%).

Using the recorded traces, we then analysed the conformational dynamics of four HJ constructs: the reference HJ with both dyes covalently attached; the two HJs where one dye is supplied by r-labels whilst the other one is attached covalently; and the complete REFRESH-HJ.

We used the E traces (see *Methods*) to generate FRET efficiency frequency distributions and dwell-time histograms for the four different HJs (Figure 5 and S4 for the single r-labels). All structures showed similar FRET distributions, with peaks at E values of ≈ 0.25 and ≈ 0.75. The relative abundance of the two fractions for the HJ (reference: low FRET: ≈ 70 % high FRET: ≈ 30 %, REFRESH: low FRET: ≈ 51 %, high FRET: ≈ 49 %) indicate an equilibrium constant of K_high→low_ ≈ 2.3 for the reference HJ, and a K_high→low_ ≈ 1.0 for the REFRESHFRET HJ. The relative abundance indicates that the low-FRET state is energetically slightly favoured in the reference HJ, however, the difference between the states is ~ 2-fold smaller when using r-labels. Notably, the shift in the equilibrium when only one dye is supplied as an r-label is much stronger for the red one (K_high->low_ ≈ 1.2) than for the green one (K_high->low_ ≈ 2.0), suggesting that the majority of the equilibrium shift is induced by the attachment of the red docking strand and r-label (see Figure S4).

**Figure 3.**
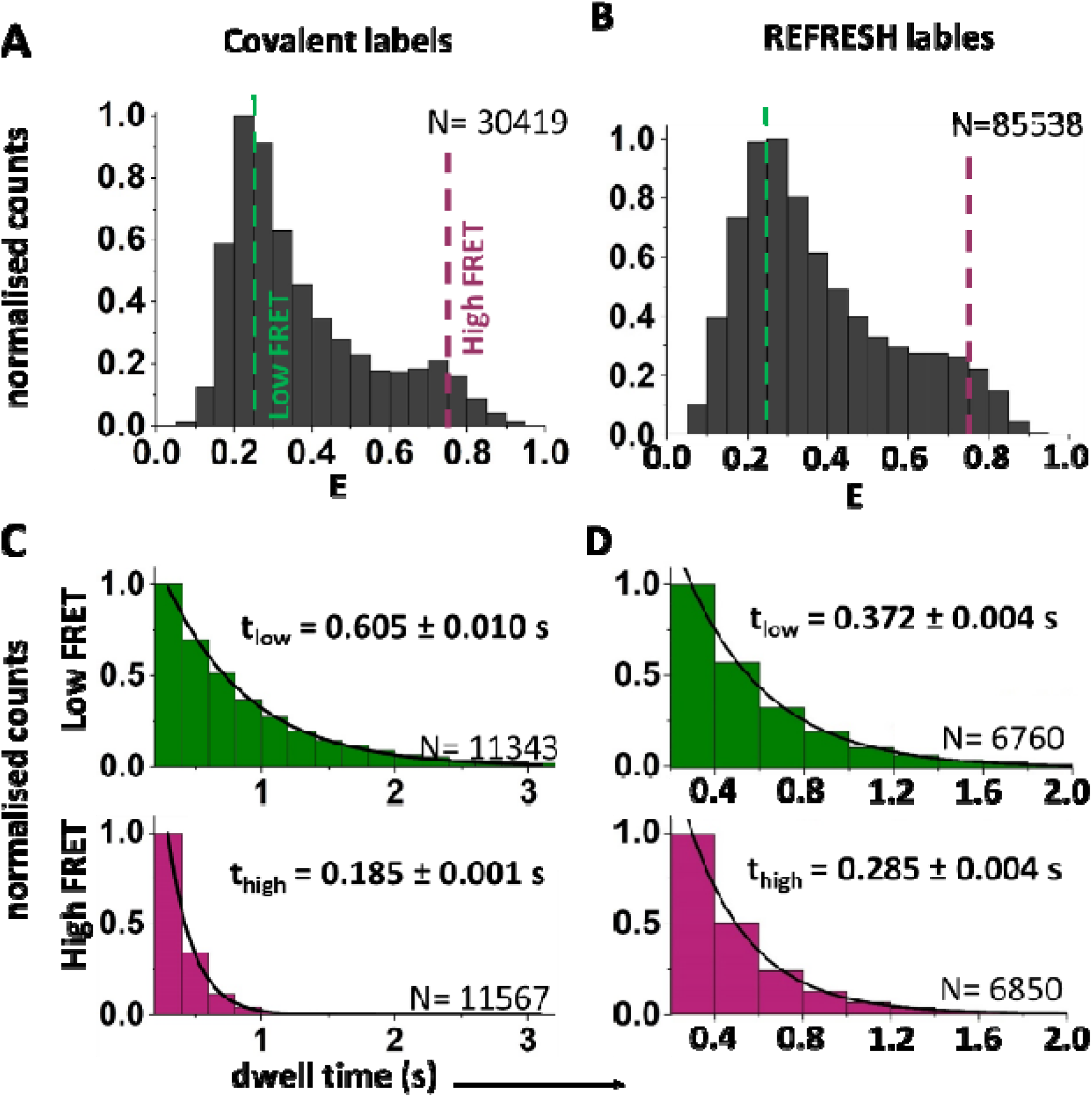
Monitoring conformational dynamics using REFRESH-FRET: Representative data from one experiment. A-B. FRET histograms of the reference HJ with covalent labels (A) and the HJ with exchanging donor and acceptor (B). C–D. Dwell time histograms of the high and low FRET states for the reference HJ (panel C; data from 64 molecules), and the HJ with exchanging acceptor label (D; data from 101 molecules), the errors stated are fitting errors. N, number of frames.

We also determined the kinetic constants for the interconversion of FRET states. In the REFRESH-HJ, at 100 nM red and 300 nM green r-label, the interconversion rates were k_high→low_ = 3.45 ± 0.04 s^−1^ and k_low→high_ = 2.69 ± 0.03 s^−1^, equating to a K_high->low_ ≈ 1.3. In comparison, the reference HJ showed a k_high→low_ = 5.51 ± 0.22 s^−1^, k_low→high_ = 1.61 ± 0.04 s^−1^, and a K_high→low_ ≈ 3.4. These values agree well with the values obtained from the population fitting and further show that, the shift in equilibrium when comparing reference and REFRESH HJ is due to a change in the stability of both states. An interconversion constant K_high→low_ near unity is consistent with previously published literature on the HJ (Gilboa et al reported a K_high→low_ of ≈ 3.7, while McKinney et al reported a K_high→low_ of ≈ 1 across different [MgCl_2_] ^[36,38]^).

Our results clearly establish that we can use REFRESH-FRET to accurately resolve conformational dynamics well below the second timescale over observation times on the scale of hours, spanning five orders of magnitude.

Ultimately, the observation time span is limited by the survival time of the target molecule, especially the docking strand. In DNA-PAINT, photo-destruction of docking strands has been reported; however, DNA-PAINT uses 10-25-fold higher laser powers than in our experiments.^[39]^ Additionally, DNA-PAINT experiments are often performed without photostabilisation, which both preserves fluorescent dyes and prevents damage to DNA structures (such as docking sites or r-labels).^[19]^ Consistent with this, we have observed only a few traces (<5%) which permanently enter a dark state after some time (or show significant reduced on-times).

## Conclusion

In most single-molecule fluorescence studies, photobleaching severely limits the available photon budget from a reporter fluorophore, with each measurements needing to carefully balance temporal resolution, overall observation span, and photons collected per frame, with the latter feature determining the spatial or FRET resolution of the measurement).

By circumventing this limitation, REFRESH allows monitoring processes at high temporal resolution over long observation spans, opening many new opportunities for singlemolecule studies. Most prominently, this allows access to long-lived or rare states in slow reactions, which would be mis-characterised in a bleaching-limited system. Such states and transitions have been suggested for many proteins, nucleic acids and their complexes and might be critical to understanding mechanisms and defining rate-limiting steps.

Further, REFRESH enables observations on the same molecule throughout multiple rounds if its activity, e.g., the same protein molecules can be monitored during several rounds of the catalysed reaction of the same or different substrate, or in the presence of other molecules which may alter the protein behaviour. Complementary to this idea, we can also monitor a substrate molecule through several rounds of processing by different enzymes, e.g., we envision systems that detect the repeated synthesis and/or degradation of specific RNA molecules both *in vitro* and *in vivo* using transient fluorescent in situ hybridization (FISH).^[40–43]^Importantly, the reversible nature of renewable labelling would permit interactions with RNA-processing proteins without the interference caused by stably bound FISH probes. We also envision of large range of other potential applications, such as the development of improved biosensors, and new cellular imaging methods.

## Supporting information

SI_Kummerlin et al, 2022

## Acknowledgements

We thank ChristofHepp for critical reading of the manuscript, Martin Rieu for the help with the statistical analysis of the dwell time data, and the MICRON Advanced Bioimaging Facility (supported by Wellcome Strategic Awards 091911/B/10/Z and 107457/Z/15/Z) for access to a Nanoimager microscope in their facilities. This work was supported by the Wellcome Trust (110164/Z/15/Z to A.N.K.), the UK BBSRC (BB/R008655/1 to A.N.K.), a UK EPSRC studentship (project 2440758 to M.K.), and a doctoral fellowship of the BoehringerIngelheimFonds (to M.K.). The illustrations in Figures 1, 2A, and 3A-B were created using BioRender.com

## Notes

### Competing Interest Statement

The work was performed using miniaturized
commercial microscopes from Oxford Nanoimaging, a company in which Achillefs N Kapanidis is a co-founder and shareholder.

### Summary of Updates

revised title

