## Supplementary material for "Bleaching-resistant, near-continuous single-molecule fluorescence and FRET based on fluorogenic and transient DNA binding": SI_Kummerlin et al, 2022

### **MATERIAL & METHODS**

#### **Holliday Junction annealing and immobilisation on surfaces.** Oligos were obtained from Metabion and Merck, dissolved to a final concentration of 100 μM, and stored at -20ºC (for Sequences, see Table S1). HJ components (strands: HJ-H, HJ-B, HJ-X or HJ-XI, HJ-R or HJ-RI) were mixed in annealing buffer (200 mM Tris–HCl pH 8.0, 500 mM NaCl, 1 mM EDTA) at 2-4 μM, and then annealed in a thermocycler (program: heating to 90°C, then cooling to 25°C at 2°C/min, storing at 4°C).

HJs were immobilised via a biotinylated H-strand binding NeutrAvidin on coverslips coated by polyethylene-glycol (PEG). In wells of silicone gaskets, 20 µl of the HJs (100 -500 pM) were incubated for 10-30 s, followed by washing three times with 200 µl PBS. Subsequently, 30 µl of DNA imaging buffer (200 mM MgCl_2_, 10 mM NaCl, 50 mM HEPES pH 7.4, 6 mM BSA, 3 mM TROLOX, 1% Glucose, 40 µg/ml catalase and 0.1 mg/ml glucose oxidase) containing the stated r-label concentrations were added. For hour-long acquisition of FRET dynamics, the buffer was renewed at a rate of 6 μL/min, allowing for a complete volume exchange every 5 min.

**Movie Acquisition:** Single-molecule fluorescence movies were collected using the Nanoimager-S single-molecule fluorescence microscope (Oxford Nanoimaging). The microscope was used as a widefield single-molecule fluorescence microscope with objective-based total internal reflection fluorescence (TIRF) illumination mode, with the excitation angle set at 53.6°. We performed the imaging using continuous-wave excitation (532 nm for Cy3B and 640 nm for ATTO647N) or alternating laser excitation (ALEX) mode, with the laser powers of 12-13% (2.9-3.6 mW) at 532 nm; and 6% (1.4 mW) at 640 nm. In all experiments, we used 100-ms exposures and ALEX movies were recorded at 100ms/frame/excitation, leading to traces with 200-ms temporal resolution.

**Fluorescence Trace extraction:** Movies were corrected for lateral drift as follows: localisations were found using Picasso^[1]^ ‘localize’ and were then loaded in ‘render’ and un-drifted by redundant cross-correlation (RCC). The created drift file was used in a custom MATLAB (MathWorks) script to un-drift individual frames, which were then combined using FIJI.^[2]^

For all data except the 1-hr trace, fluorescence intensity vs. time traces were extracted and background-corrected using TwoTone.^[3]^ The program extracts the fluorescence intensity in the green and red channel upon green excitation (DD and DA, respectively), and in the red channel upon red excitation (AA). Traces were manually inspected and any traces in which multiple molecules were detected, were discarded.

For one-colour experiments, the DD signal was used for localisation, and the AA signal was plotted as intensity vs time traces. In FRET experiments, all three signals were used to calculate the apparent FRET efficiency E and donor-acceptor stoichiometry S were calculated as follows:^[4]^

E =DA/(DD+DA)	(1)

S= (DA + DD)/(DD+DA+AA)	(2)

Two-dimensional E-S plots were used to select data points which contain both donor and acceptor dyes, for which E histograms were plotted.

Further analysis and HMM fitting were performed using ebFRET.^[5]^ We fitted two distinct FRET states and extracted dwell time histograms for each state. By fitting these with a single exponential, the transition rates were determined. Stated rates with errors are means and standard deviations, respectively, of three independent experiments. The figures show exemplary data from one of multiple experiment per condition.

All traces and histograms were plotted using Origin (OriginLab).

**Determining binding-kinetics of the r-labels.** For extraction of dwell times from our fluorescence time traces, we performed HMM fitting on the intensity vs time traces obtained for both r-labels. Fitting worked well on the green traces, but for the red label, the multiple emission states of ATTO647N combined with the two intensity levels resulting from two bound fluorophores lead to difficulties for HMM fitting. We thus resulted to using localisation-based information extracted using Picasso^[1]^ for fitting dwell times for the red r-label. Localisations were extracted using the ‘localize’ tool and then un-drifted using redundant cross correlation (RCC). All remaining localisations were filtered for in sx/sy for the main population and linked allowing one dark frame between localisations. From this data, the length of binding events (=on times) and the dark time between events (=off time) can be extracted.

To give readers straightforward numbers for the hybridisation kinetics in the main text, we have calculated mean values of t_off_ and t_on_, and also the inverse values as k_on_ and k_off_, respectively.

From previous studies, we know, however, that the hybridisation kinetics are indeed better described by a bi-exponential decay with two independent decay constants. To further characterise the binding behaviour, we thus fitted a bi-exponential distribution to the dwell times of both r-labels using a maximum-likelihood-estimator in Matlab (MEMLET^[6]^). The fitted function takes into account the minimum measurable dwell time (one frame) and the discrete nature of the observable values. Confidence intervals for fitted decay times were determined via bootstrapping (Table S2). The results show that the green label at 100 nM has on-rates of k_on,1_ =0.31 s^‑1^ and k_on,2_ =0.031 s^‑1^ with 75.8% of events following k_on,1_. The off-rate constant are k_off,1_ =0.48 s^-1^ and k_off,2_ =0.038 s^‑1^ with 34.4% of all events following k_off,1_ (see Figure S1A, C). For the red r-label, on-rates of k_on,1_ =3.50 s^‑1^ and k_on,1_ =0.12 s^‑1^ with 90.5% described by k_on,1_ and off-rates of k_off,1_ =0.90 s^-1^ (78.4%) and k_off,1_ =0.031 s^-1^ were observed at 20 nM (Figure S1B,D).

**Characterising self-quenching R-label strands.** We additionally tested a construct terminally labelled with two ATTO655, two ATTO647N and Dabcyl, and ATTO647N and BHQ3. All constructs were 11-nt long, except the ATTO655 labelled probe, which had a length of 8 nt.

To characterise the level of quenching, we performed ensemble measurements, assessing the absorption spectra and the fluorescence of the quenched r-labels in absence and presence of up to 100-fold excess of complementary DNA.

The fluorescence spectra were measured at a scanning spectrofluorometer (PTI) using 1-s integration time per 1-nm-wavelength intervals using 100 μL r-labels in buffer (50 mM HEPES, pH 7.4; 200 mM MgCl_2_, 10 mM NaCl, 0.1 % BSA, the same buffer as used for single molecule measurements) to a final concentration of 100 nM. Complementary DNA was added stepwise to achieve different concentrations until saturation of the signal was observed (0.1-10 μM). Samples were excited at 520 nm (containing Cy3B), 620 nm (containing ATTO647N) or 640 nm (containing ATTO655).

The emission spectra (Figure S2) were recorded in presence of 0 to 10 µM complementary DNA (complementary DNA was added until the fluorescence signal saturated). Upon hybridisation, the increased stiffness of dsDNA forces the dye-dye interactions apart and thus should de-quench the probes. All red r-labels (Figure S2A-D) show an increase in fluorescence upon addition of an increasing concentration of complementary DNA, most prominent the 2xATTO647N and the ATTO647N-BHQ3 probes (4-fold and 6-fold increase, respectively). The level of fluorescence in the ATTO647N-BHQ3 probe is, however, still lower than the fully quenched 2xATTO647N. We reasoned that there is significant FRET from ATTO647N to the BHQ3 even in the hybridised state, which prevents fluorescence emission and thus renders the probe unsuitable for our purpose. For further single-molecule experiments, we selected the 2xATTO647N r-label as a red label, since it proved to be the most suitable out of the tested selection.

For the green r-label, we observe a 16-fold increase in fluorescence intensity under the same conditions (Figure S2E), which suggests that indeed increasing the distance between dark quencher and fluorophore in the de-quenched state (compared to the ATTO647N-BHQ3 pair) is allowing for strong emission once hybridised with complementary DNA.

**Table S1**: Sequences of DNA strands. The HJ is formed by nucleotides in capitals, other nucleotides are involved in r-label binding.

| Strand name | Sequence | Modification |
| --- | --- | --- |
| HJ-B | 5′-CCCTAGCAAGCCGCTGCTACGG | -- |
| HJ-H | 5′-CCGTAGCAGCGAGAGCGGTGGG | 5’-biotin |
| HJ-R | 5′-CCCACCGCTCTTCTCAACTGGG | 5’-ATTO647N |
| HJ-X | 5′-CCCAGTTGAGAGCTTGATAGGG | 5’-Cy3B |
| HJ-RI | 5′-aa aaa ggg aaa-CCCACCGCTCTTCTCAACTGGG | -- |
| HJ-XI | 5′- ttc aac att tct tct CCCAGTTGAGAGCTTGATAGGG | -- |
| rI-2xATTO647N | 5’-ttt ccc ttt tt | 5’ and 3’ ATTO647N |
| rI-ATTO647N-Dabcyl | 5’-ttt ccc ttt tt | 5’ ATTO647N, 3’Dabcyl |
| rI-ATTO647N-BHQ3 | 5’-ttt ccc ttt tt | 5’ ATTO647N, 3’ BHQ3 |
| comprI | 5’-aa aaa ggg aaa | -- |
| gI-Cy3B-BHQ2 | 5’-aga agt aat gtg gaa | 5’ Cy3B, 3’ BHQ2 |
| compgI | 5′- ttc aac att tct tct | -- |
| 8mer-2xATTO655 | 5’-tcc acc gt | 5’ and 3’ ATTO655 |
| Comp-8mer | 5’-ac ggt gga | -- |

| label | state | parameter | mean | lower 95% CI | upper 95% CI |
| --- | --- | --- | --- | --- | --- |
| green | off | A | 0.757 | 0.752 | 0.761 |
|  |  | t1 (in s) | 3.189 | 3.151 | 3.227 |
|  |  | t2 (in s) | 31.080 | 30.616 | 31.544 |
|  | on | A | 0.341 | 0.337 | 0.345 |
|  |  | t1 (in s) | 1.491 | 1.455 | 1.528 |
|  |  | t2 (in s) | 24.907 | 24.706 | 25.108 |
| red | off | A | 0.905 | 0.905 | 0.906 |
|  |  | t1 (in s) | 0.285 | 0.284 | 0.285 |
|  |  | t2 (in s) | 8.051 | 8.005 | 8.098 |
|  | on | A | 0.783 | 0.783 | 0.784 |
|  |  | t1 (in s) | 1.105 | 1.101 | 1.108 |
|  |  | t2 (in s) | 31.708 | 31.571 | 31.845 |

**Table S2**: Parameter mean and 95% CI from bootstrapping (200 iterations) the binding kinetics of the r-labels.

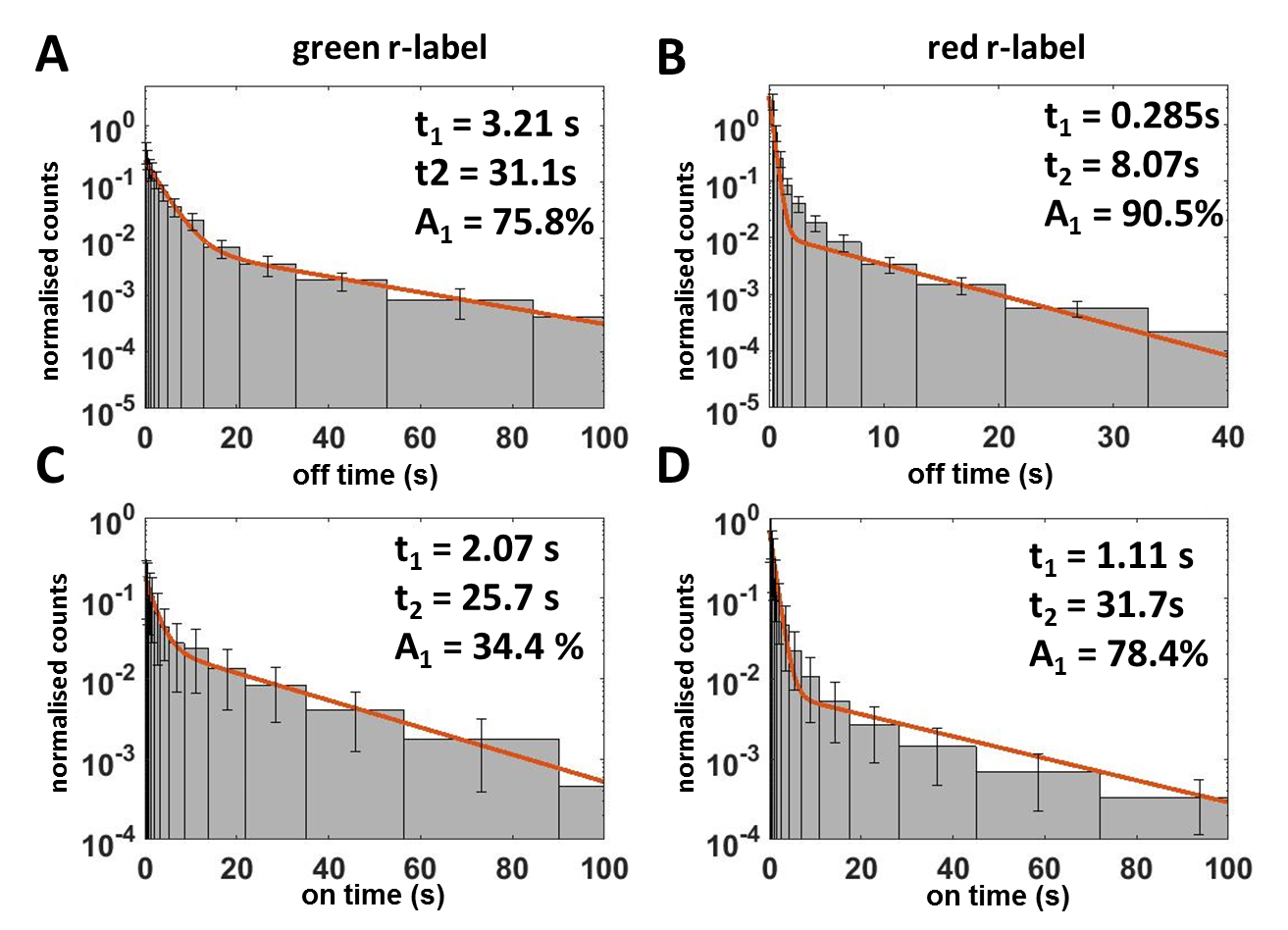

**Figure S1**: Characterisation of the binding kinetics for the r-labels. The data was fitted to a bi-exponential distribution using a MLE algorithm in Matlab (MEMLET^[6]^, red graph). For display purposes, the data were binned with increasing bin size, counts were normalised by the bin width. Error bars on bins are derived from bootstrapping (200 iterations).

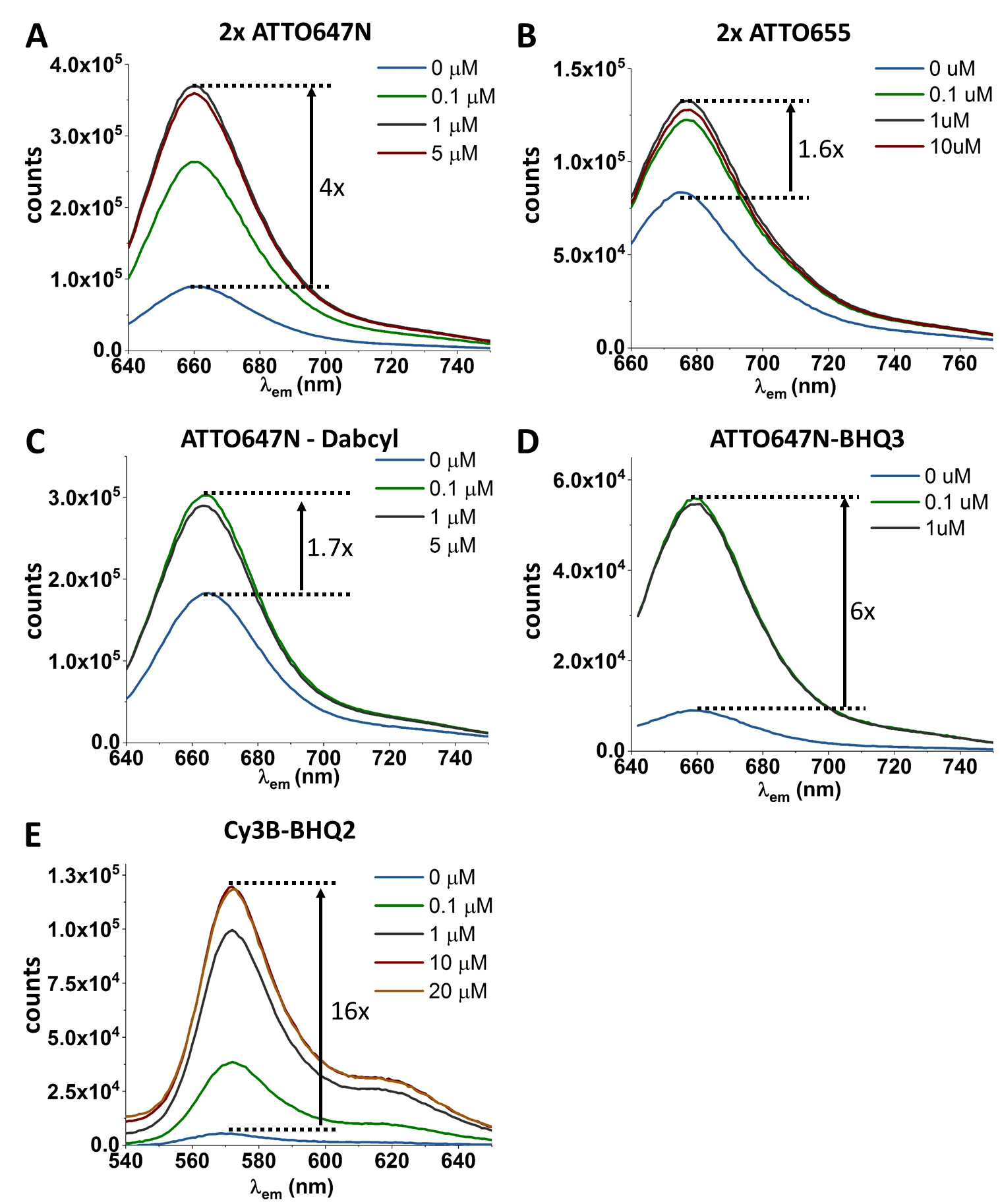

**Figure S2**: Characterisation of quenching efficiency in double-labelled r-label strands. Fluorescence spectra of probes terminally labelled with two ATTO647N (**A**), two ATTO655 (**B**), ATTO647N and Dabcyl (**C**), ATTO647N and BHQ3 (**D**), or Cy3B and BHQ2 (**E**) at various concentrations of complementary ssDNA.

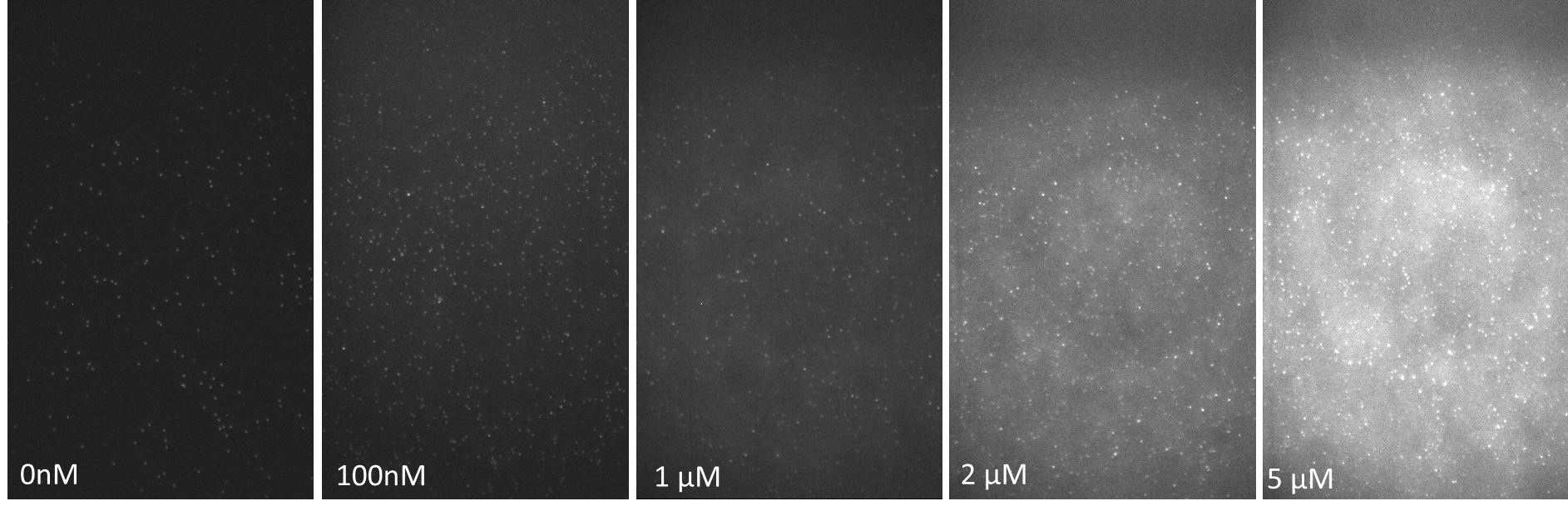

**Figure S3:** Individual frames imaged at various concentrations of green r-labels (0-5μM) in the imaging buffer. At 0 nM, the observed spots come from immobilised Cy3B. The subsequence images show spots of green r-labels at various concentrations binding to complementary, immobilised docking strands. Through the fluorogenic nature of the r-lables, individual targets can still clearly be identified at 5μM of r-lables in the imaging buffer.

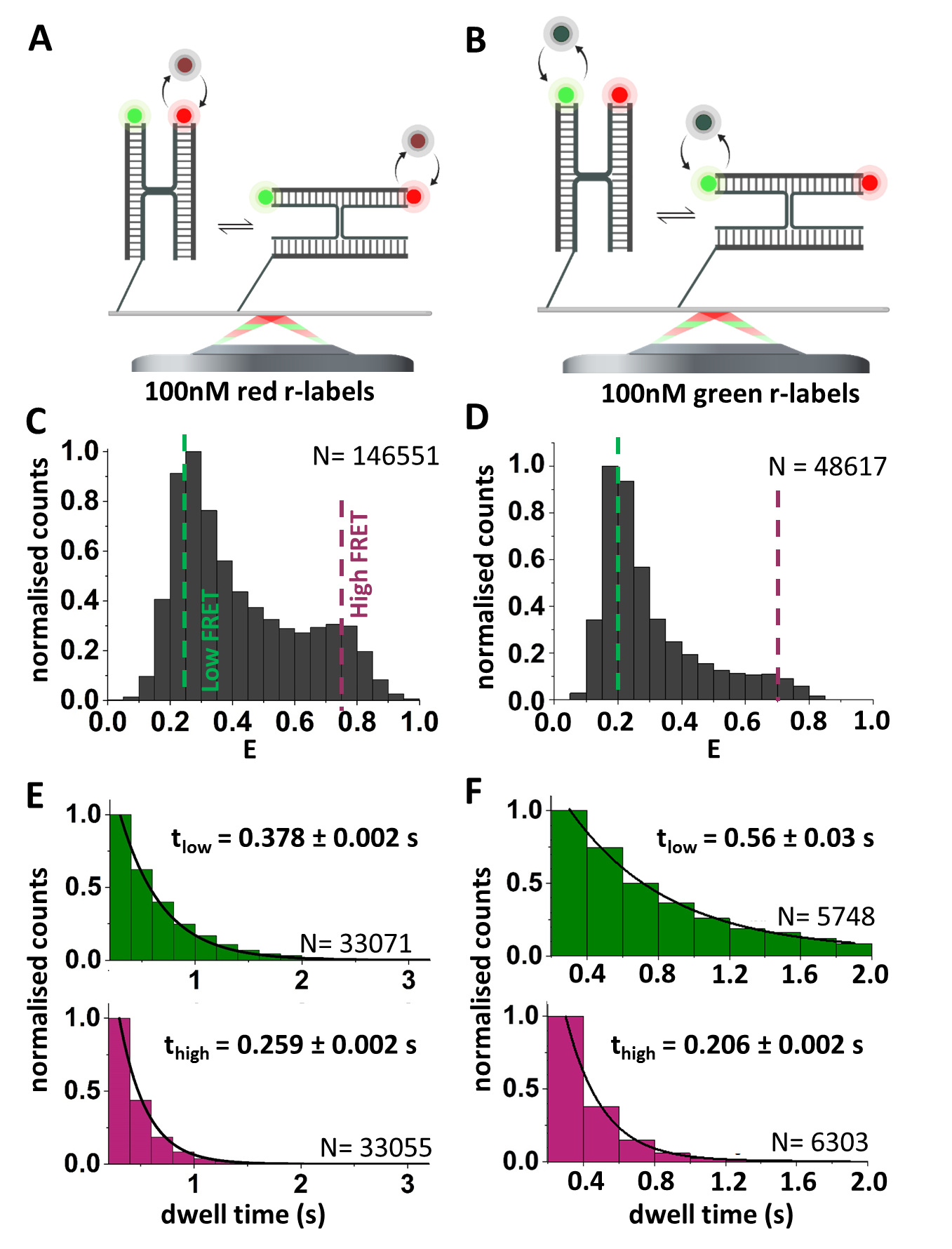

**Figure S4**: Conformational dynamics in the HJ observed with single r-labels. FRET distributions (**C-D**) and dwell time histograms (**E-F**) for the HJ with only (**A, C, E**) or green (**B, D, F**) r-label, as indicated by the schematics in **A** and **B**, respectively. Data from one representative experiment of 179 (red r-label) and 72 molecules (green r-label).

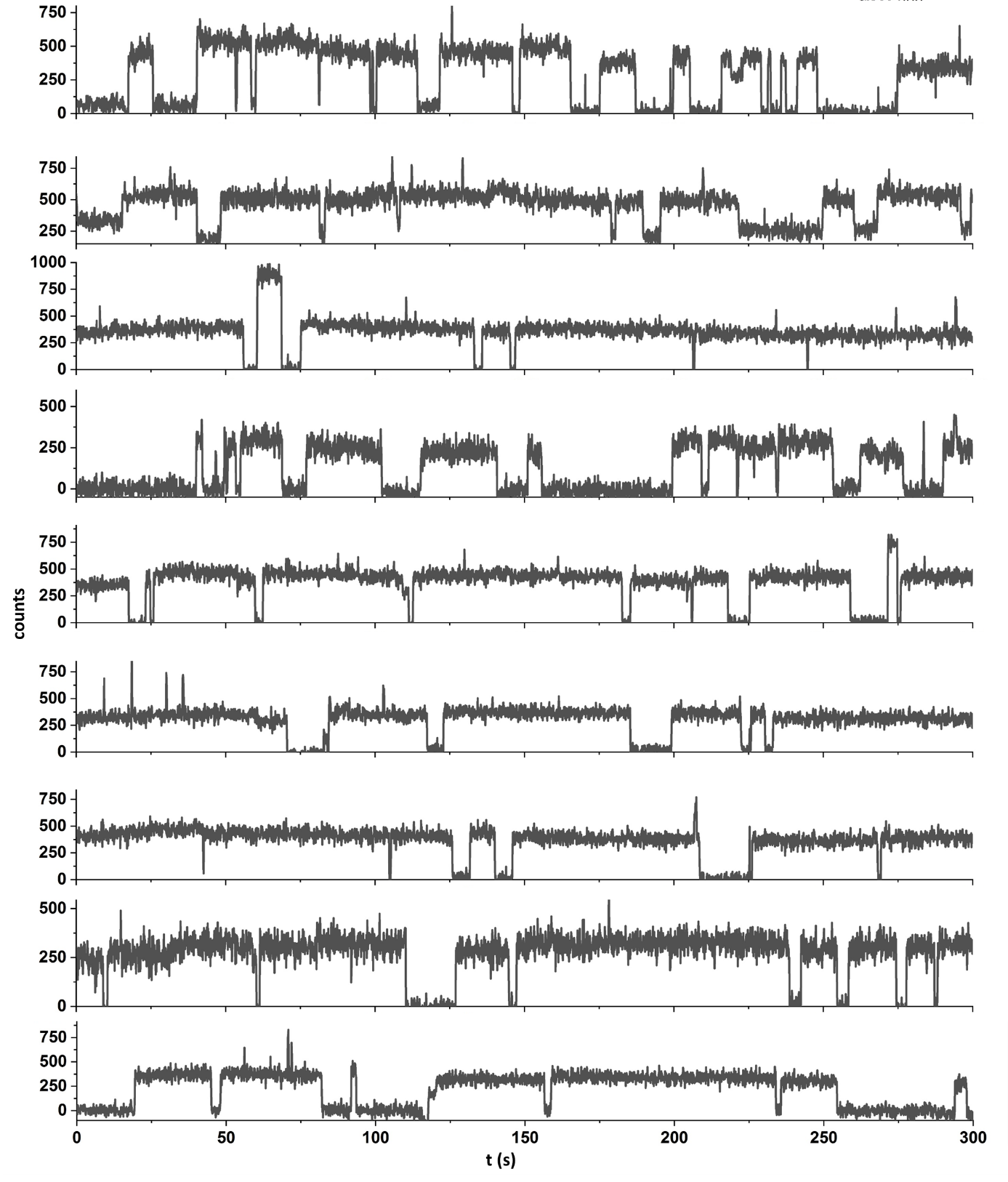

**Figure S5:** Additional traces of the green r-label binding at 100 nM. Repeated binding can be observed by the signal rising to ≈450 counts. Occasionally, the signal reaches higher counts (≈ 800 counts), which indicates intervals of Cy3B fluorescence without functional BHQ2.

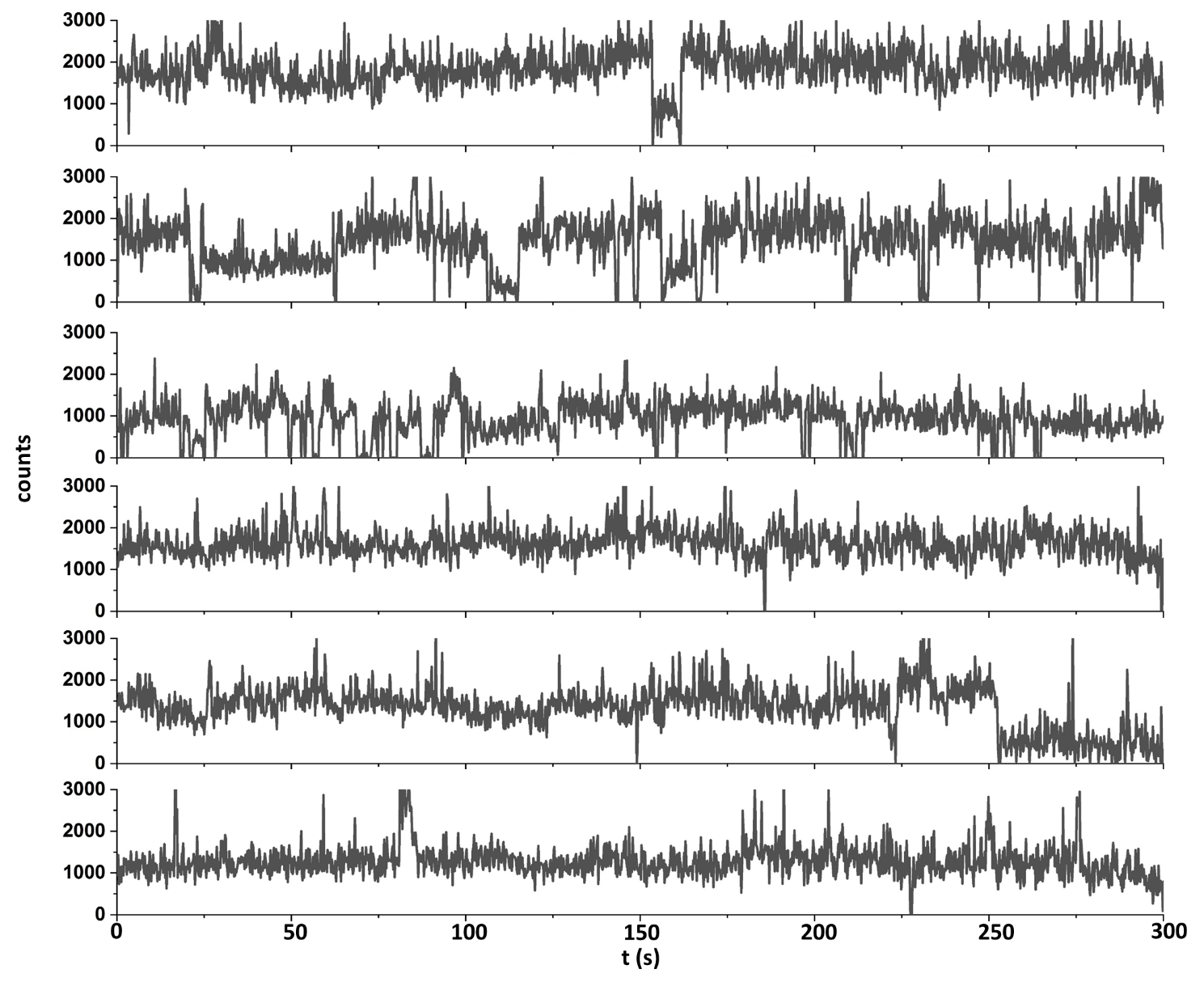

**Figure S6:** Additional traces of the red r-label binding (100 nM). Repeated binding can be observed by the signal rising to ≈1000 - 1500 counts. Occasionally, the signal shows lower counts (≈ 750 counts), which indicates intervals of only one functional ATTO647N dye emitting.

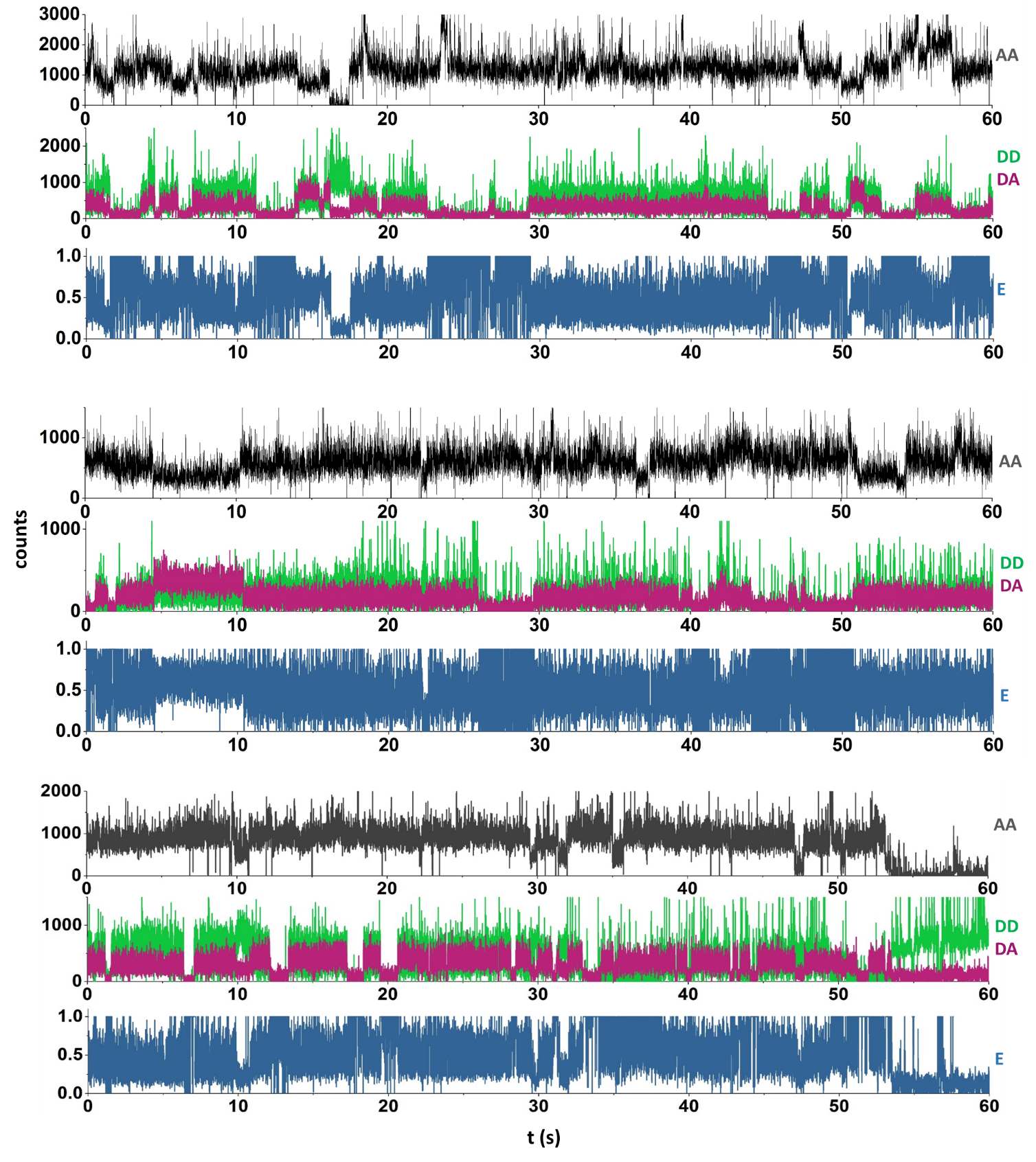

**Figure S7:** Additional traces of the green (300 nM) and red r-label (100 nM) binding in the FRET regime on the HJ with AA (top, grey), DD, and DA channel (middle, green and magenta, respectively) and calculated FRET efficiency (E, bottom, blue). The anti-correlated fluctuations in the DD and DA channels (middle) and the fluctuating E trace (bottom) indicate a dynamic interchange between a high-FRET state (E ≈ 0.75) and a low-FRET state (E ≈ 0.25).
